# MiR-126 Improves the Therapeutic Effect of EPC in Hypertensive Ischemic Stroke

**DOI:** 10.1101/2025.03.27.645845

**Authors:** Jun Huang, Bing Zhao, Xiaodong Li, Liqiang Dingli, Mengxia Fu, Yaying Song, Meijie Qu, Dongrui Chen, Guo-Yuan Yang, Pingjin Gao

**Affiliations:** Department of Cardiovascular Medicine, Department of Hypertension, State Key Laboratoryof Medical Genomics, Shanghai Key Laboratory of Hypertension, Shanghai Institute of Hypertension, Ruijin Hospital, Shanghai Jiao Tong University School of Medicine; Emergency Department, Ruijin Hospital, School of Medicine, Shanghai Jiao Tong University, Shanghai, China; Renji Hospital, School of Medicine, Shanghai Jiao Tong University; Neuroscience and Neuroengineering Research Center, Med-X Research Institute and School of Biomedical Engineering, Shanghai Jiao Tong University, Shanghai, China

**Keywords:** endothelial progenitor cells, exosome, ischemic stroke, miR-126, spontaneously hypertensive rats

## Abstract

**Background and Purpose:** Hypertension promotes circulatory endothelial inflammation, reduces endothelial progenitor cell (EPC) function, and impairs their therapeutic potential following ischemic stroke. MiR-126 is known to regulate vascular development and angiogenesis. This study investigates the therapeutic potential of miR-126 by overexpressing it in EPCs to enhance their efficacy in hypertensive stroke conditions.

**Methods:** Adult spontaneously hypertensive rats (SHRs, n=118) underwent permanent suture middle cerebral artery occlusion (MCAO) to induce ischemic stroke. One week post-stroke, animals were injected with EPCs overexpressing miR-126 or control EPCs. Treatment effects were evaluated over 35 days, using neurological scoring, infarct volume measurements, behavioral testing, and assessments of neuroinflammation, blood pressure, and angiogenesis. Exosomes from miR-126 overexpressing EPCs were isolated and analyzed for mechanistic studies.

**Results:** *In vitro*, miR-126 enhanced EPC angiogenic function under stress. *In vivo*, miR-126 modified EPC treatment improved functional recovery, reduced infarct volume, and promoted angiogenesis compared to controls in SHRs. Furthermore, miR-126 treatment preserved the blood-brain barrier, reduced peripheral immune cell infiltration, and modulated neuroinflammation. Exosomes derived from miR-126 overexpression EPCs also promoted angiogenesis and reduced endothelial activation under stress conditions.

**Conclusions:** EPC overexpressing miR-126 provides neuroprotection by enhancing angiogenesis, reducing ischemic injury, and preserving blood-brain barrier integrity in hypertensive stroke models. This approach modulates neuroinflammation and improves neurological outcomes, suggesting gene-modified EPCs as a promising strategy for ischemic stroke therapy, particularly under hypertensive conditions.

## Introduction

Chronic hypertension is a major risk factor for ischemic stroke and adversely affect neurological recovery after stroke^1^, ^2^. Despite its prevalence, the mechanisms through which hypertension contributes to stroke pathology and impairs rehabilitation are not fully understood. Hypertension is known to promote inflammation in the brain endothelium, often referred to as cerebral vascular inflammation, and results in a significant loss of stem cell function, especially endothelial progenitor cells (EPCs), which are essential for vascular repair after stroke. Impaired EPC function and reduced numbers of these cells in the bone marrow are common sequelae of hypertension and stroke, further hindering recovery^3-8^.

EPCs play a key role in cerebral angiogenesis within the ischemia area, which is crucial for neurovascular unit repair and the restoration of neurological function after ischemic stroke^9^, ^10^. Although EPC based treatment have shown promise in preclinical studies^11^, ^12^, their efficacy remains limited, especially under hypertension conditions. Due to inflammatory mediators (such as cytokines and alarmins), and the hypoxia status in the peri-infarct region, all of these factors impair the EPC function and disrupt vascular repair, reducing treatment effectiveness^8^, ^13^. Thus, understanding the mechanisms that enhance EPC survival and angiogenesis function under hypertensive and ischemic conditions is critical for improving stroke recovery outcomes.

MiR-126-3p (miR-126) is a highly expressed microRNA in endothelial cells, has been shown to promote vascular remodeling and angiogenesis^14^, ^15^. Studies indicate that increased levels of miR-126 are associated with blood brain barrier (BBB) protection and improved outcomes in brain ischemia^16^, ^17^. However, its expression is downregulated under the hypertensive or ischemic conditions^18^. In this study, we hypothesized that overexpressing miR-126 in EPCs could enhance their angiogenic function and improve recovery after stroke in hypertensive rats. Additionally, we explored the role of exosomes secreted by miR-126-modified EPCs as a potential mechanism for enhancing angiogenesis and modulating inflammation in the brain.

Our results demonstrate that miR-126 overexpression in EPCs improves long-term neurological functional recovery in hypertensive stroke models, with exosome-mediated effects contributing to enhanced angiogenesis and reduced inflammation.

## Materials and Methods

### Isolation and culture of EPCs

Human umbilical cord blood was obtained from the International Peace Maternity and Child Health Hospital of China (IPMCH, Shanghai, China). The procedure for EPC isolation and purification was performed as described previously^19^. Briefly, monocytes were isolated using Ficoll separation. EPCs (5–8 passages) were used in the subsequent experiments. EPC were identified using immunofluorescence staining and flow cytometry as described previously^20^.

### Flow cytometry assay

Single cells suspensions were labeled with fluorescein isothiocyanate (FITC)-conjugated mouse anti-human CD31, FITC-conjugated mouse anti human CD133, phycoerythrin (PE)-conjugated mouse anti-human CD34, PE-conjugated mouse anti-human CD90, and PE-conjugated mouse anti-human KDR (all from BD Biosciences). The sample for the negative control was treated with a non-specific antibody that shared the same isotype as the specific antibody used in the experiment. Flow cytometry was performed on a BD Biosciences FACScan. Flow cytometric data were analyzed using FlowJo (Ashland, OR) software.

### Middle cerebral artery occlusion in SHRs

Animal studies were reported in accordance with Animal Research: Reporting of In Vivo Experiments 2.0 (ARRIVE) guidelines. Procedure for using laboratory animals was approved by the Institutional Animal Care and Use Committee (IACUC), Ruijin Hospital, Shanghai Jiao Tong University, Shanghai, China. Adult male SHRs, weighing 300–350 grams were used in this study. MCAO was performed according to the previously described methods with minor modifications^21^. Briefly, SHRs were anesthetized with ketamine/xylazine (100 mg/10 mg/kg; Sigma, St. Louis, MO). Maintaining the body temperature 37±0.3°C using a heating pad. Isolation of the left external and internal carotid artery, a silicone-coated suture (RWD Life Science, China) was gently inserted from the external carotid artery stump to the internal carotid artery, and stopped at the opening of the middle cerebral artery. Laser Doppler flowmetry (Moor Instruments, Devon, UK) was used to test the occlusion.

### Plasma or oxygen-glucose deprivation (OGD) culture condition

Plasma was collected 3 days after MCAO or sham surgery in SHRs^22^. SHR stroke or sham plasma was used for media conditioning in EPC culture. For OGD condition, cells were maintained with deoxygenated glucose-free ECM (Gibco) in a chamber containing an anaerobic gas mixture (95% N_2_ and 5% CO2) at 37 °C for 4 h. Then, the cultures were removed from the chamber, and the OGD supernatant was replaced with a maintenance medium. For the standard (normoxic) condition, cells were maintained in serum under an ambient atmosphere at 37 °C.

### RNA isolation and miRNA expression analysis

Total RNA was isolated using miRNeasy kit (Qiagen, Hilden, Germany) according to the manufacturer’s instructions. The qPCR of mature miRNA was performed using the miRCURY LNA Universal RT miRNA PCR kit consisting of SYBR Green Master Mix (Exiqon, Woburn, MA) with LNA primers for hsa-miR-126-3p and hsa-miR-126-5p. *U6* was used to normalize the expression of the target gene.

### miRNA transfection

Synthetically derived mature-sequence miR-126-mimic molecules (Sheng Gong, Shanghai, China) were transfected in EPC at a pre-optimized concentration (40 nmol/L) to overexpress miR-126 and repress the target expression. Random sequence miR-mimic was used as a negative scramble control (40 nmol/L, Sheng Gong). miR-126 or control mimic molecules were transfected into EPC with Lipofectamine 3000 48 h before serum conditioning, according to manufacturer instructions.

### Tube Formation Assay

Tube formation assays were based on a previously described procedure^23^. Briefly, 36 h after transfection of miRNA mimics, EPCs were stimulated under OGD or SHR stroke plasma and seeded on Matrigel^TM^ (BD Biosciences, Franklin Lakes, NJ) in a 96-well plate. Tube lengths were measured and quantified using National Institutes of Health Image*J* software (Bethesda, MD).

### Migration Assay

Migration assays were based on a previously described procedure ^23^. Briefly, transwell Boyden chambers (BD Biosciences, Franklin Lakes, NJ) were inserted into 24-well plates, 500 μl of normal medium was added to the lower compartment. EPCs (1×10^4^ cells/well) transfected of miRNA mimics were added to the upper chamber. After stimulation in OGD or add SHR stroke plasma for 6 h, the cells that migrated to the lower membrane surface were stained with hematoxylin and counted manually under a microscope. Five random fields from each well were selected, and three wells in each group were examined.

### Proliferation Assay

Proliferation assays were based on a previously described procedure ^24^. Briefly, 1×10^4^ cells/well were seeded in 96-well plates with 100 μl of medium/well. Cell Counting Kit-SF (Dojindo Laboratories, Kumamoto, Japan) was used to perform a WST-8 ([2-(2-methoxy-4-nitrophenyl)-3-(4-nitrophenyl)-5-(2,4-disulfophenyl)-2H-tetrazolium]) assay. Cell Counting Kit-8 reagent was added to the cell culture medium to a final concentration of 5 μl/100 μl and incubated for an additional 4 h at 37°C. The absorbance was measured at 450 nm against a reference wavelength of 630 nm on a microplate reader (ELx800; Bio-Tek Instruments, Winooski, VT).

### Bromodeoxyuridine (BrdU) administration

BrdU (Sigma) was solubilized in normal saline at a concentration of 10 mg/mL injected intraperitoneally at a dosage of 50 mg/kg twice each day from 21–35 days after stroke or sham surgery for 2 weeks before the animals were sacrificed on day 35.

### Measurement of infarct volume

Brain infarct volume was determined by crystal violet staining (Sigma). Brains were frozen in −40°C isopentane and 20 μm thick coronal sections were sliced from the anterior commissure to the hippocampus. 20 consecutive sections were mounted on slides and stained by crystal violet. The infarct size was quantified using NIH Image*J* software, and the infarct volume was calculated as described previously^21^.

### Neurological deficiency assessment

The neurological assessment was performed before cerebral ischemia and on days 7, 14, 21, 28, and 35 after MCAO. Briefly, the neurological deficiency assessed by an investigator blinded to treatment and according to a modified neurological severity score (NSS) 0–15 grading system (normal score, 0; maximum deficit score)^25^. Among the scores of injuries, 1 score point is awarded for the inability to perform the test or for the lack of a tested reflex; thus, the higher the score, the more the injury. The test correlated with infarct volume and has been described previously^26^, ^27^.

### Exosome isolation

Exosomes were isolated and purified as described previously^28^. Briefly, after EPC^miR-126^ was washed twice with phosphate-buffered saline (PBS), cells were switched to an exosome-free cell culture medium to culture for 48 h. The supernatant was collected and clarified by sequential ultracentrifugation at 2000 *g* for 30 min, 10000 *g* for 30 min, and 100000 *g* for 70 min at 4°C. Then, the exosomes were washed once with PBS at 100000 *g* for 70 min and resuspended in PBS for further identification. The exosomes characterization was identified by a transmission electron microscope (TEM, Thermo Scientific, Waltham, MA). Exosomes markers (CD9 and CD63) and tumor susceptibility gene 101 (*TSG101*) were detected by Western blot analysis. The PKH26 was labeled according to the instruction manual (Sigma).

### In Vivo murine model

Exosomes were isolated from EPC^miR-126^ and co-cultured with HUVECs for 48 h, then HUVECs were resuspended in phenol red-free Matrigel (BD Biosciences). Cell/Matrigel mixture (1.5×10^6^ cells /200 ml) was injected subcutaneously into the backs of 6-week-old male athymic nu/nu mice (n=4 per group). The animals weight gain and matrigel plug size were monitored at day 7 and day 14. On the day 14, the animals were euthanized and matrigel plugs were harvested. The matrigel plugs were frozen at −80°C, 20 μm thick sections were sliced for hematoxylin-eosin (HE) staining.

### Immunostaining Quantification

Immunofluorescence was performed on 20 thick serial brain sections. The procedure is as previously described. Information of primary antibodies is described in **Suppl Table 1**. For MPO immunohistochemistry, after stained with MPO antibody, followed by incubation with biotin-conjugated anti-rabbit antibody and ABC regents. DAB Peroxidase Substrate (Sigma) were used to developed the sections. For BrdU staining, after washed three times in TBST, brain sections were transferred to 95°C for 10min in 10mM sodium citrate, then washed three times in TBST. Sections were incubated in 3M HCl for 30 min at 37°C, blocked for 1.5h in TBS++, and then incubated in primary antibody for 72 h. After incubate with secondary antibody, sections were mounted with Flouromount. Fluorescence images were captured under a confocal microscope (Leica Microsystems, Wetzlar, Germany).

### Statistical Analysis

All data are presented as mean ±standard error of the mean (SEM). The comparisons between the two groups were carried out using unpaired two-tailed t-tests. One-way analysis of variance (ANOVA) with Tukey’s multiple comparisons test was used for multiple-parameter analysis. Statistical analysis was performed using GraphPad Prism 8 (GraphPad Software, San Diego, CA).

## Results

### miR-126 expression in EPC is decreased under stress conditions

Our investigation into miR-126 expression in hypertension revealed that circulating miR-126 levels are downregulated in hypertensive individuals compared to those with normal blood pressure adults. Additionally, analysis of previously published data showed that EPC function was inversely correlated with blood pressure and is impaired under hypertensive conditions. To further explore the miR-126 expression in EPC under stress condition, we first assessed the expression levels of miR-126 (miR-126-3p) and miR-126’ (miR-126-5p) in EPC under normal conditions. EPCs isolated from human umbilical cord blood were characterized (**Suppl Fig. 1A-C**), and both miR-126 and miR-126’ levels were found to be higher in EPCs compared to those in the HUVECs (**Fig. 1A**).

**Fig. 1.**
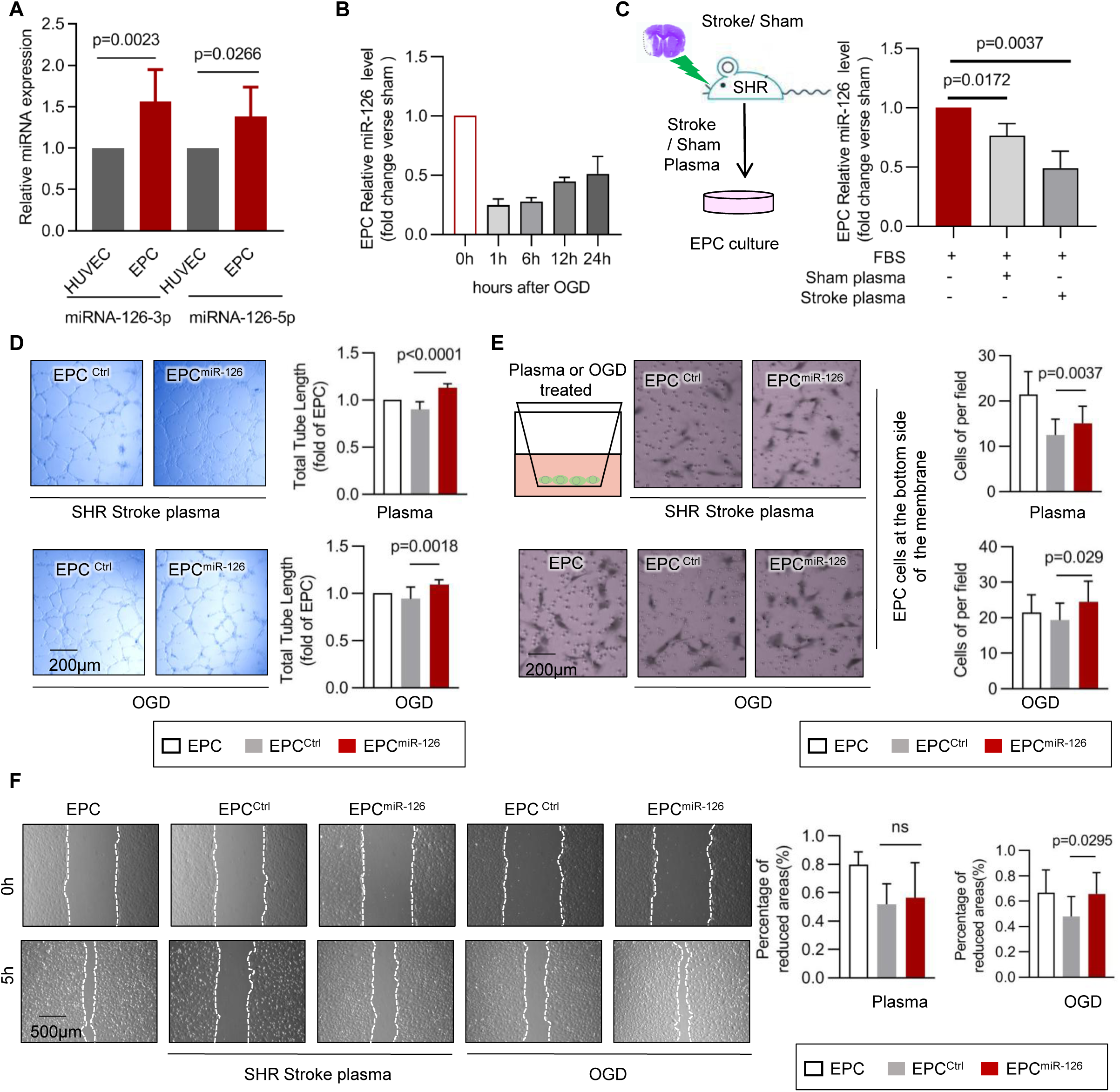
MiR-126 enhances EPC function under hypertensive brain ischemia stress conditions. A) miR-126-3p (miR-126) and miR-126-5p expression in EPCs and HUVECs. **B**) miR-126 expressing of EPCs under OGD stimulation. **C**) Schematic illustration of experiments: plasma was collected from 24 h post-stroke or sham SHR, and used for conditioning media in EPC cultures. miR-126 expressed in EPC was detected after being incubated with stroke or sham plasma. **D**) Tubule formation of EPC^ctrl^ and EPC^miR-126^ after stroke plasma or OGD incubation. **E**) Migration of EPC^ctrl^ and EPC^miR-126^ in transwell cultures showing assay design (upper left), hematoxylin-stained images (middle), cell counts (right). **F**) Scratch test showing of EPC^ctrl^ and EPC^miR-126^ migration after stroke plasma or OGD incubation.

Following ischemic stroke in the context of hypertension, EPCs are exposed to two major stressors: 1) the inflammatory plasma environment induced by hypertension and stroke, and 2) the hypoxia conditions in the peri-infarction area. To further explore the expression of miR-126 in EPC under stress conditions, we used an in vitro culture system to simulate these stress conditions. EPCs were exposed to plasma from SHR stroke model or subjected to an oxygen-glucose deprivation (OGD) to mimic the ischemic environment. Our results showed that miR-126 expression in EPC is significantly downregulated when exposure to SHR stroke plasma compared to sham plasma (**Fig. 1C**), as well as under OGD conditions (**Fig. 1B**). These findings suggested that the soluble factors present in hypertension plasma after stroke, along with hypoxic conditions in the peri-ischemic region, contribute to the decrease expression of miR-126 in EPCs, potentially impairing their angiogenic and repair functions.

### Overexpression of miR-126 enhances EPC function under stress conditions *in vitro*

To investigate the role of miR-126, we overexpressed miR-126 mimics or mimic control in EPCs. After confirming that these agents were successfully taken up by EPCs(**Suppl Fig. 1D**), we evaluated the overexpression of miR-126 on EPC tube formation, migration, and survival under two stress conditions: SHR stroke plasma or OGD. Notably, exposure to SHR stroke plasma enhanced EPC tube formation and transwell migration with miR-126 overexpression, although scratch migration was unaffected. In contrast, under OGD conditions, miR-126 improved EPC tube formation, transwell migration, and scratch migration (**Figs. 1D–F**). However, miR-126 did not significantly increase EPC survival under either stress condition(**Suppl Fig. 1E**), suggesting that miR-126 promotes EPC tube formation and migration rather than proliferation. These findings imply that hypertension, stroke plasma, and peri-ischemic hypoxia impair EPC migration and tube formation, and that miR-126 overexpression can mitigate these disfunction.

### EPC ^miR-126^ enhanced functional recovery in SHR after MCAO

To evaluate in vivo effects of miR-126-modified EPC, we transduced EPC with miR-126 using lentiviral vector (EPC^miR-126^) and used GFP-transduced EPCs as a control (EPC^GFP^, **Suppl Fig. 2A-B**). The expression of miR-126 was over 20-fold higher in EPC^miR-126^ compared to EPC^GFP^ (**Suppl Fig. 2C**).

As shown in **Fig. 2A**, SHRs were pre-trained 7 days before MCAO surgery. On day 7 post-MCAO, EPC^miR-126^ (10^6^ cells) or control EPC^GFP^ were resuspended in normal saline and injected via tail vein for 3 consecutive days, bypassing the acute inflammation phase. We next assess examined the impact of EPC^miR-126^ on the functional recovery after MCAO. As shown in **Fig. 2F**, EPC^miR-126^-treated exhibited significantly improved NSS performance on days 28 and 35 post-MCAO compared to both the EPC^GFP^ or NS group. In addition, performance on the adhesive removal, bean balance, and rotarod test showed significant improvements in the EPC^miR-126^ group relative to the controls (**Fig. 2B**). To further evaluate the therapeutic effect of EPC^miR-126^ in the later stage of MCAO under hypertensive conditions, we assessed ischemic brain volume on day 35 post-MCAO (**Fig. 2D**). The results indicated that EPC^miR-126^ group had significant smaller brain ischemic volumes compared to the EPC^GFP^ or NS groups (**Fig. 2G**).

**Fig. 2.**
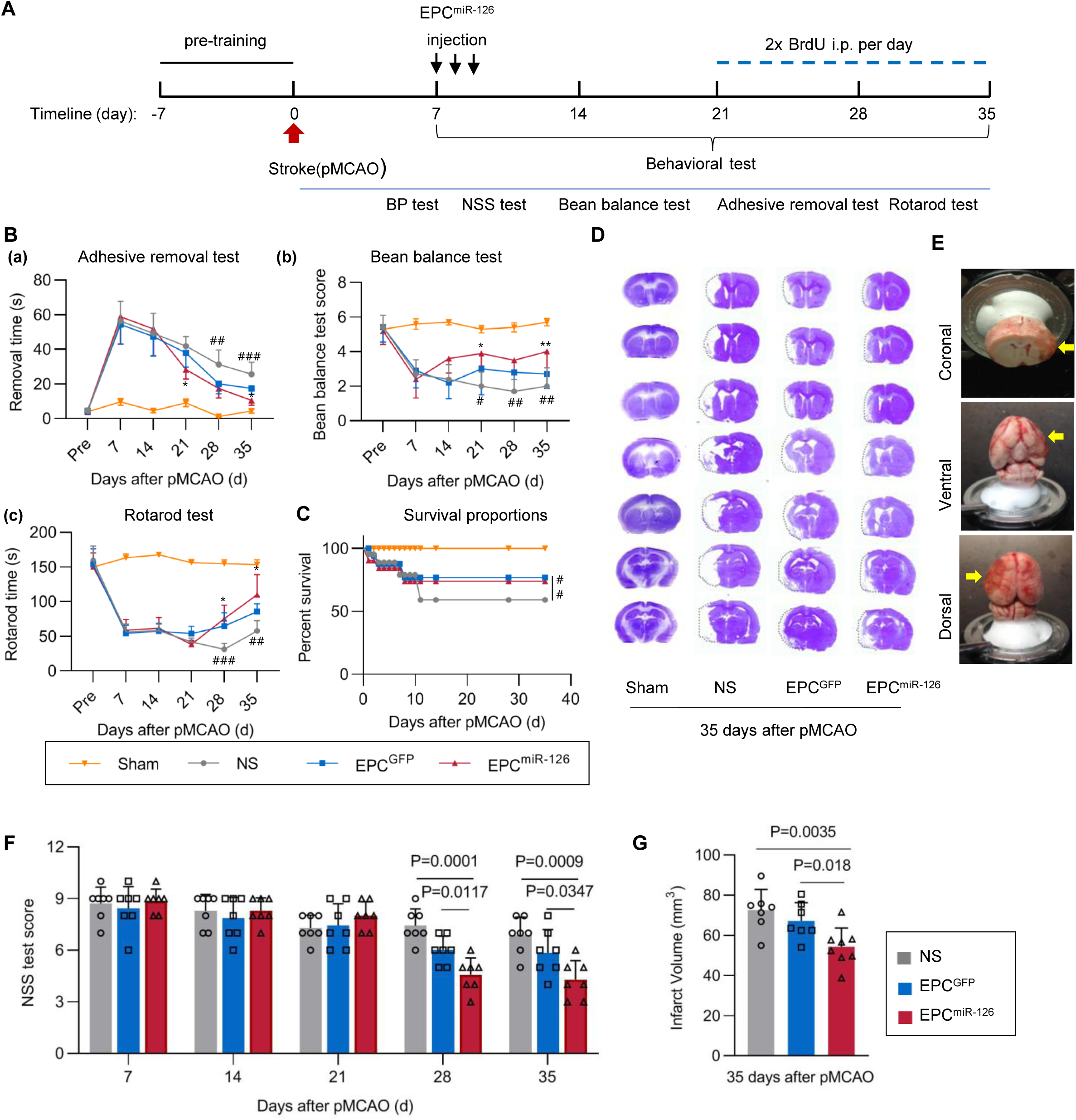
EPC^miR-126^ reduced infarction size, improve survival, and enhances neurological recovery after MCAO in SHR rats. **A**) Schematic of experimental design and time course, including EPC administration, behavioral test, BrdU injection. **B**) Adhesive removal test (**a**), bean balance test (**b**), rotarod test (**c**) were detected at different time points after stroke. N=10 per group. **C**) Post-stroke mortality rates were lower in EPC^miR-126^ group and EPC^GFP^ group. Log-rank test. N=12–16 per group. \**P<0.05, **P<0.01, ***P<0.001,* EPC^miR-126^ vs EPC^GFP^; #*P<0.05, ##P<0.01, ###P<0.001,* EPC^miR-126^ vs NS; 2-way repeated-measures ANOVA followed by Holm-Sidak post hoc multiple comparisons test were used. **D**) Cresyl violet stained coronal sections of SHR brain at 35 days after MCAO. **E**) Photos show coronal, ventral and dorsal of the brain ischemic area. Arrows indicate the ischemic area. **F**) Neurological scores of different treated groups after MCAO. Data are mean±SEM, N=6-8 per group. **G**) Quantification of the infarct volumes. Data are mean±SEM, N=6–7 per group.

Together, these data demonstrated that treatment with EPC^miR-126^, starting 7 day after SHR stroke, enhanced long-term neurological functions and sensory motor recovery. However, no significant differences were observed in systolic or diastolic blood pressure or body weight between the groups (**Suppl Fig. 3A-B**).

### EPC^miR-126^ promoted angiogenesis in SHR after MCAO

Next, we investigated the angiogenic effects of EPC^miR-126^ in SHR after MCAO. Bromodeoxyuridine (BrdU) was injected on days 21to 35 post-MCAO to label newly generated cells. As shown in **Fig. 3A**, the number of CD31^+^/BrdU^+^ cells in the peri-infarct area (both in the peri-infarct cortex and stratum) was significantly higher in all the therapeutic groups compared to the NS or sham group. On day 35 post-MCAO, the EPC^miR-126^ group exhibited the greatest increase in CD31^+^/BrdU^+^ cells compared to the EPC^GFP^, NS, and sham group (**Fig. 3A-B**).

**Fig. 3.**
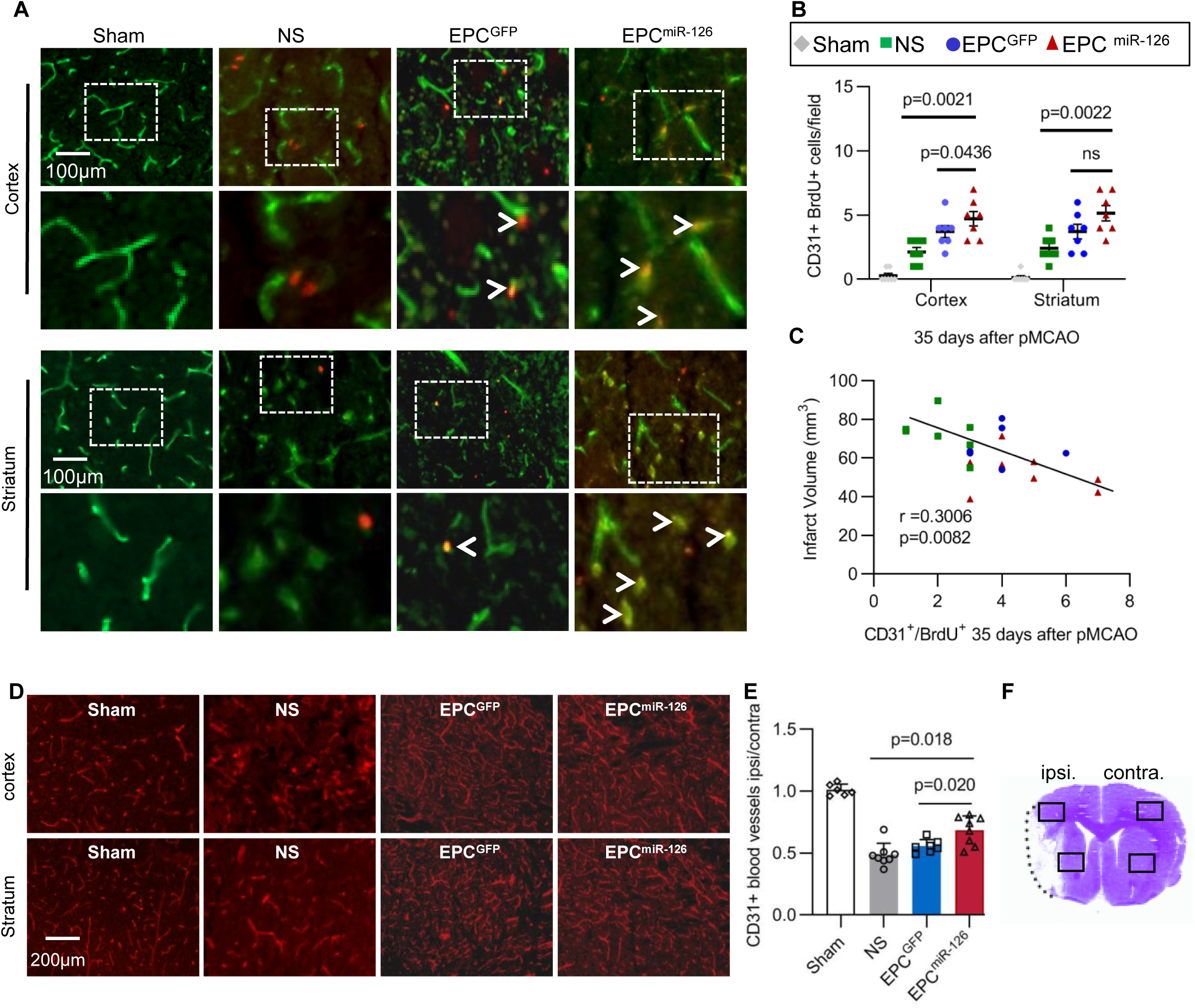
EPC^miR-126^ rescues angiogenic function in SHR rats following MCAO. **A**) Photograph of CD31 and BrdU double staining of ipsilateral hemisphere at 5 weeks after MCAO. The arrowheads point to CD31^+^/BrdU^+^ colocalized newly formed microvessels. **B**) Quantification of the number of BrdU^+^ and CD31^+^ double-immunostained cells. N=6–10 per group. **C**) Pearson correlation between the number of BrdU^+^ and CD31^+^ and infarct volume in the ipsilateral hemisphere at 5 weeks after MCAO. **D**) Representative photograph of CD31^+^ microvessel in the sham, NS, EPC^GFP^ and EPC^miR-126^ treated group in the ischemic perifocal region of cortex and striatum at 5 weeks after MCAO. **E**) Bar graphs showing the ratio of number of microvessels in the ipsilateral/contralateral of the ischemic brain 5 weeks after MCAO. Data are mean±SEM, N=6–10 per group. ipsi.=ipsilateral, contra.=contralateral

To explore whether angiogenesis was linked to a reduction in infarct size, we analyzed the relationship between the number of newly generated CD31^+^/BrdU^+^ cells and the infarct volume. A negative correlation was observed between these two variables, indicating that increased angiogenesis was associated with smaller infarct volumes (**Fig. 3C**). Additionally, the density of CD31^+^ blood vessels was significantly higher in the peri-infarct region of all therapeutic groups compared to the NS groups, with the highest density observed in the EPC^miR-126^ group (**Fig. 3D**). Furthermore, the ratio of blood vessels density in the ipsilateral to the contralateral hemisphere was greater in the EPC^miR-126^ group compared to all other groups (**Fig. 3E-F**). These results suggested that the transplantation of EPC^miR-126^ promotes angiogenesis and enhances blood vessel intensity in the late stage of SHR stroke brain, contributing to improved tissue repair and recovery.

### EPC^miR-126^ protected BBB integrity and reduced brain immune cell infiltration and glial activation

To investigate the role of EPC^miR-126^ in ischemic stroke recovery in SHR, we assessed the stability of BBB tight junction ZO-1 and occludin Immunofluorescence-staining for ZO-1 and occludin on day 35 post-MCAO revealed continuous ZO-1 or occludin expression in the EC layer of brain vessels in the sham group (**Fig. 4A**). In the NS group, MCAO caused significant disrupted of the microvessel walls with discontinuous ZO-1or occludin labeling. In contrast, EPC^miR-126^ treated mice showed smoother and more continuous ZO-1 and occludin staining along the microvessel walls (**Fig. 4A**, arrowheads). Quantification of total gap length showed less disruption in the EPC^miR-126^-treated group post-MCAO compared to the NS and EPC^GFP^ groups (**Fig. 4A**).

**Fig. 4.**
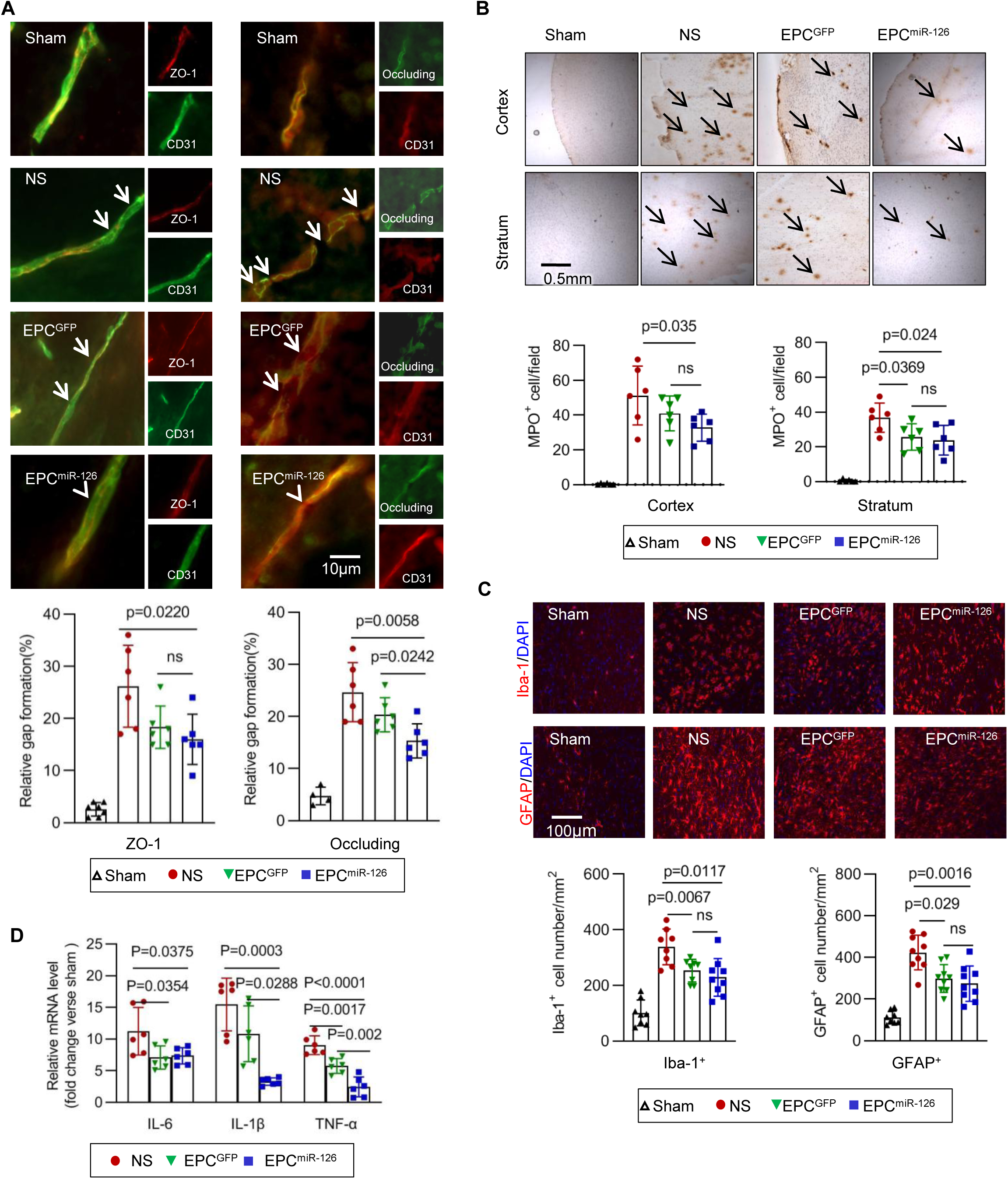
EPC^miR-126^ protects BBB and reduced immune cell infiltration and glial activation after MCAO. **A**) Immunofluorescence staining of ZO-1 and occludin expression in the sham, NS, EPC^GFP^ and EPC^miR-126^ treated SHR post-stroke. Microvessels in the sham group showed a continuous and linear labeling of tight junction protein ZO-1 or occluding along the CD31^+^ vessels. ZO-1 or occludin showed a discontinuous, irregular distribution in microvessels in NS treated group. Less disruption and fewer gaps (arrowheads) were detected in the EPC^GFP^ or EPC^miR-126^ treated group (arrowheads). Bar graph shows the quantification of gap length. Data are mean±SEM. N=6 animals per group. **B**) MPO staining in sham, NS, EPC^GFP^ and EPC^miR-126^ treated SHR brain after MCAO. Bar graph shows the quantification of MPO cells in the cortex of stratum in different treated groups. Data are mean±SEM. N=6 animals per group. **C**) Immunoreactivity for microglial marker Iba-1 and astrocytic marker GFAP was assessed under different treatment groups at day 35 post-stroke. Reactivity was notably reduced with EPC^miR-126^ treatment. Bar graph shows the quantification of Iba-1^+^ or GFAP^+^ cells. Data are mean±SEM, N=6 animals per group. **D**) EPC^miR-126^ Treatment inhibited the brain mRNA expression of inflammatory cytokines in stroke SHR. N=6 animals per group.

To assess peripheral leukocyte infiltration, we measured myeloperoxidase (MPO) levels, which indicate neutrophil accumulation, on day 3 after treatment. The number of MPO^+^ cells was significantly higher in the NS group and lower in the EPC^miR-126^ and EPC^GFP^ treatment groups with no significant difference between the latter two (**Fig. 4B**). We also examined glial activation and neuroinflammation in the ischemic brain. Both EPC^miR-126^ and EPC^GFP^ treatment significantly reduced the number of reactive astrocytes and microglia compared to the NS group (**Fig. 4C**). Furthermore, mRNA expression levels of proinflammatory cytokines IL-6, IL-1β, and TNF-α was significantly lower in the EPC^miR-126^ group (**Fig. 4D**). These results indicated that EPC^miR-126^ treatment protected BBB integrity, reduced neutrophils infiltration into the ipsilateral hemisphere, and attenuates neuroinflammatory response following SHR MCAO.

### Exosomes mediated miR-126 trafficking from EPC to EC

While angiogenesis was detected in the peri-infarct area, inflammation in the brain was reduced, and neurobehavioral improvements were noted following EPC^miR-126^ treatment, the injected EPC^miR-126^ or EPC^GFP^ cells was rarely detected on day 35 after MCAO in SHRs. These findings suggested that, aside from directly participating in the ischemic area, the injected cells may also contribute to neurovascular repair through paracrine or endocrine mechanisms, particularly via the release of exosomes that carry bioactive molecules. To explore this, we isolated exosomes from EPC^miR-126^ by ultracentrifugation, and electron microscopy revealed that the exosomes were within the size range as 40–150 nm (**Fig. 5A**). Western blotting analysis confirmed the presence of exosome-specific markers (CD63, TSG101) and exosome-associated HSP70 protein (**Suppl Fig. 4A**).

**Fig. 5.**
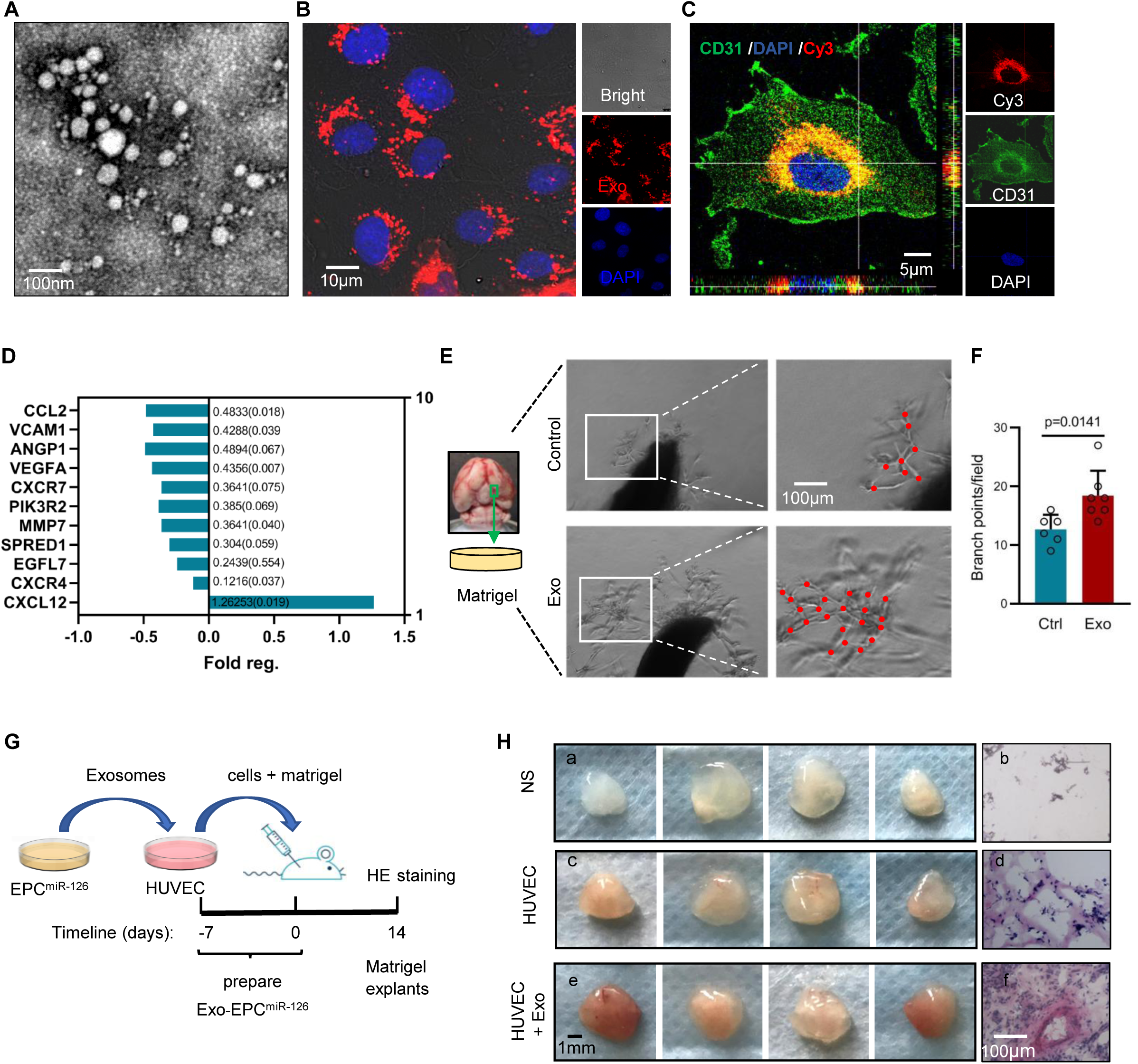
Exosomes derived from EPC^miR-126^ enhance angiogenesis *in vivo* and *in vitro*. **A**) Electron micrograph of exosomes isolated from EPC^miR-126^. **B**) Immunofluorescence staining of HUVEC DAPI and PKH26-labelled EPC^miR-126^ exosomes after 24 h incubation. **C**) Confocal microscopy analysis of HUVEC after incubation with miR-126 Cy3 labelled exosomes for 24 h. **D**) Fold change and adjusted P values of targets of miR-126 in HUVEC transcription after 3 days incubation with EPC^miR-126^ exosomes. **E**) Exosomes from EPC^miR-126^ increased angiogenic sprouts of middle cerebral artery explants. **F**) Bar graph shows quantification of angiogenic branch points of middle cerebral artery explants (n=6 per group). Data are shown as mean±SEM. **G**) Experimental design and time course of the in vivo matrigel explants experiment. **H**) Explants removed 14 days after implantation of matrigel seeded with NS, HUVEC and HUVEC with exosomes from EPC^miR-126^ (4 mice per group). Hematoxylin eosin staining of 14-day matrigel explants (b, d, f).

Next, we compared exosomes from EPC^miR-126^ and EPC^GFP^, and RT-PCR analysis revealed that *miR-126* levels were significantly higher in the EPC^miR-126^-derived exosome than in those from EPC^GFP^ (**Suppl Fig. 4B**). To investigate the trafficking of EPC^miR-126^ exosomes to endothelial cells, we labeled exosomes with PKH26 and co-cultured with HUVECs (**Suppl Fig. 4C**). Confocal microscopy detected labeled exosomes within HUVECs after 24 hours (**Fig. 5B**). Furthermore, to conform miR-126 transfer via exosomes, we co-cultured HUVECs with exosomes from miR-126 mimic-cy3-transfected EPC or conditional medium. After 24 hours, we detected cy3-labeled miR-126 in HUVECs (**Fig. 5C, Suppl Fig. 4D**). These results indicated that exosomes released from EPC^miR-126^ mediate intercellular miR-126 trafficking between EPCs and ECs, supporting a paracrine mechanism for neurovascular repair.

### Exosomes from EPC^miR-126^ promoted angiogenic function *in vivo* and *in vitro*

We next investigated the effects of exosomes from EPC^miR-126^ on EC function. Using polymerase chain reaction (PCR), we analyzed 11 angiogenesis-related targets of miR-126, predicted by Ingenuity Pathway Analysis (IPA, **Suppl Figs. 5A-B**), in HUVEC lysates collected 3 days after co-culture with EPC^miR-126^ exosomes. The results showed that the transcription of 10 miR-126 targets was downregulated following co-culture with EPC^miR-126^ exosomes (**Fig. 5D**). Among the downregulated targets were anti-angiogenic factors such as SPRED-1, EGFL7, MMP7, as well as CCL2, a proinflammatory cytokine that recruits proinflammatory cells to the ischemia brain after stroke^22^, ^30^, ^31^. Notably, the chemokine CXCL12, a key factor in progenitor cell recruitment, was significantly up-regulated (fold-change, 1.26; P=0.019), while its receptors, CXCR4 and CXCR7, were downregulated.

We further explored angiogenic potential of EPC^miR-126^ exosomes *in vitro* by co-cultured them with the middle cerebral artery explants from SHR in Matrigel. After 7 days, EPC^miR-126^ exosomes significantly enhanced the formation of angiogenic sprouts in MCA explants (**Figs. 5E-F**). Additionally, HUVECs were co-cultured with EPC^miR-126^ exosomes, suspended in matrigel, and injected subcutaneously into immunodeficient (nu/nu) mice (**Fig. 5G**). After 14 days, vascular networks were visible in the recipients of HUVECs co-cultured with exosomes, while only sparse vasculature was observed in the control groups (HUVECs or NS group) (**Fig. 5H**). HE staining revealed vascular rings filled with red blood cells in the exosome-treated group, but not in the control group (**Fig. 5H**).

### Exosomes from EPC^miR-126^ reduced endothelial activation post-stroke

Chemokine/chemokine receptor and integrin/adhesion molecule interactions are critical for monocyte recruitment to the endothelium following stroke. This recruitment contributes BBB disruption and endothelial activation. In our study, we observed the treatment with EPC^miR-126^ exosomes led to a decreased in the expression of chemokine receptors (CXCR4 and CXCR7) in HUVECs (**Fig. 5D**). Additionally, we found reduced mRNA expression of vcam1, a key adhesion molecule involved in monocyte recruitment (**Fig. 5D**). These findings suggested that EPC^miR-126^ exosomes may help attenuated observed endothelial activation.

To further investigate this hypothesis, we co-culture EPC^miR-126^ exosomes with HBMECs, subjecting the system to stroke- or sham-conditioned plasma, with or without GW4869. We also included OGD or normal conditions to mimic ischemic conditions (Fig. 6A). We found that stroke plasma stimulation significantly increased mRNA expression of IL-6, icam1 and vcam1 in HBMEC s co-cultured with EPC^miR-126^ exosomes. However, addition of GW4869, which inhibits exosome release, resulted in increases the expression of vcam1, and IL-6 when HBMECs were stimulated with either stroke plasma or OGD (**Fig. 6B**).

**Fig. 6.**
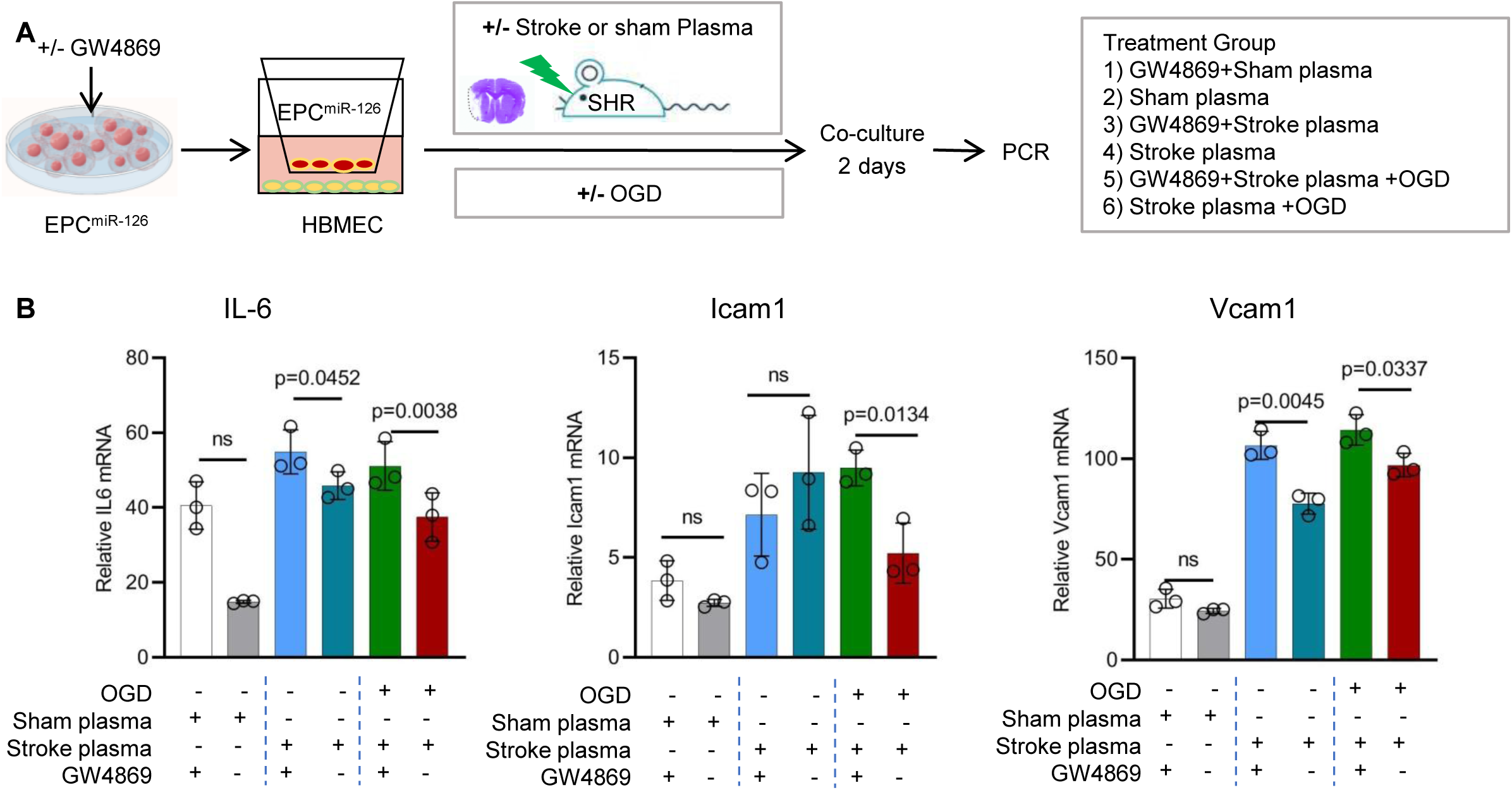
EPC^miR-126^ reduces inflammatory activation of human brain endothelium under stress conditions. **A**) Schematic illustration of experiments designs shown in (**B**). Plasma was obtained from 24 h post-stroke or sham SHR, and used for conditioning media in HBMEC cultures. **B**) Relative of IL-6, icam1 and vcam1 mRNA in HBMEC after being co-culture with EPC^miR-126^ incubated with plasma from MCAO or sham and treated with GW4869 or vehicle and treated in OGD condition or not (H test, n = 4 to 6 per group). Expression of gapdh was used as an internal control. Data are fold-change relative to HBMEC without treatment.

**Fig. 7.**
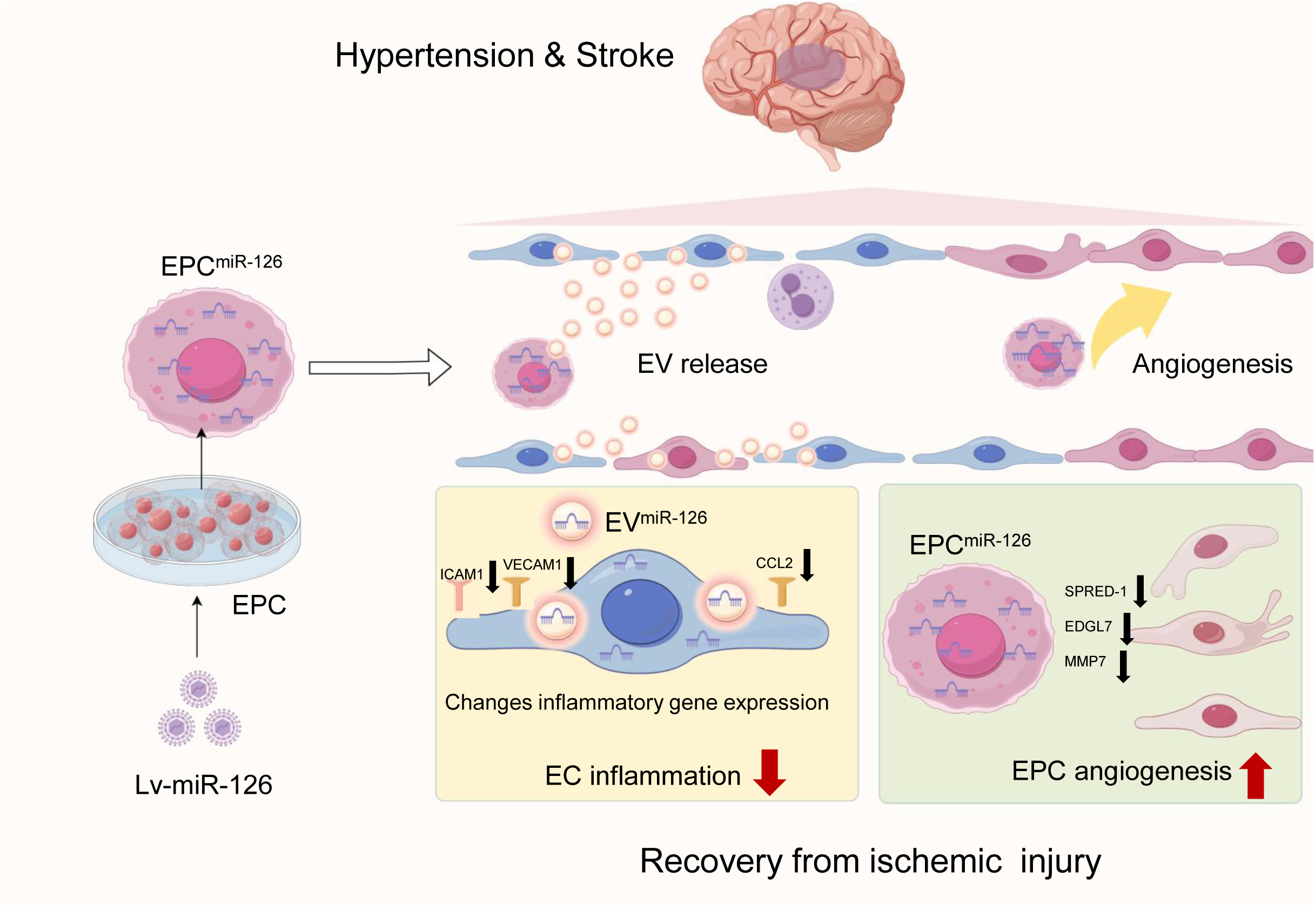
Summary of the hypothesis for EPC overexpressing miR-126 to enhance angiogenesis and reduce inflammation after focal brain ischemia in hypertensive conditions.

These results supported the hypothesis that EPC^miR-126^ exosomes reduce brain endothelial activation under hypertensive stroke conditions, potentially by modulating inflammatory pathways.

## Discussion

Hypertension is a leading cause of stroke and significantly impairs the EPC function, particularly following a stroke^32-34^. EPC are rapidly mobilized to aid in vascular repair after ischemic events. However, hypertension can negatively impair their function, especially under the compounded challenges of ischemia, hypoxia, and nutrient deficiency near the peri-infarction region^11^, ^35^. In this study, we demonstrated that miR-126 enhances the angiogenic capabilities of EPCs and promotes ischemic brain repair under hypertensions. Our findings provide new insights into how exosomes derived from EPC^miR-126^ transfer this miRNA to ECs, thereby contributing to endothelial angiogenesis, and mitigating inflammatory activation in the ischemic brain.

We detected a decrease in miR-126 levels in EPC under both the hypertensive stroke plasma stimulation and OGD conditions (**Fig. 1**). This reduction aligns with previous findings in hypertensive patients, where lower miR-126 levels were associated with impair endothelial repair function^29^, ^36^. Consistent with this, we found that overexpression of miR-126 in EPCs improved both their tube formation and transwell migration capabilities when exposure to stroke SHR plasma or OGD conditions. Although the circulating factors responsible for EPC dysfunction in hypertension or post-stroke remain to be fully identified, recent studies suggest that factors such as alarmins released from stroke necrotic brain tissue and systemic peripheral inflammation induced by hypertension may impair miRNA expression, which was in line with our findings^7^, ^22^.

Stroke induces a complex inflammatory response in the brain, characterized by an acute local sterile immune response followed by sub-acute and chronic phases, with elevated inflammatory markers persisting for more than a year in hypertensive patients^37^, ^38^. Given this, we administered EPC^miR-126^ on day 7 after SHR ischemic stroke onsets to avoid the acute inflammatory phase, provided a potential therapeutical window distinct from current stroke therapies. Our results show that EPC^miR-126^ significantly enhanced functional recovery 35 days after MCAO in SHR (**Fig. 2**). This functional improvement was associated with increased brain angiogenesis (**Fig. 3**), preserved BBB integrity, and reduced glial reactivity and peripheral immune cell infiltration (**Fig. 4**).

Consistent with prior studies, our findings indicated that stem cells based therapies can improve the stroke recovery outcomes^12^, ^39^. However, we observed that surviving of transplanted EPCs in ischemic brain was limited, with few cells retained in the tissues 4 weeks after transplantation. This suggested that, in addition to direct cell replacement, EPCs may promote vascular repair through the indirect secretion of trophic factors, forming a “biobridge” that connects EPC^miR-126^ to angiogenic sites of the ischemic brain^40^, ^41^. Among these secreted factors, exosomes play a key role in mediating angiogenesis after stroke, while the specific contribution of EPC^miR-126^-derived exosomes to vessel growth and inflammation under hypertensive and ischemic conditions remains unclear. Our study provides evidence that EPC^miR-126^-exosomes are a potent cargo for endothelial activation and angiogenesis.

We found that EPC^miR-126^-exosomes carried miR-126 and suppressed 11 targets predicted of miR-126 by IPA, most of which are known to be anti-angiogenic mRNAs (**Fig. 5**). Additionally, miR-126 was shown to promote angiogenesis via CXCL12-, CXCR7-, or MMP pathways ^42^, ^43^, while inhibiting SPRED1 and promoting VEGFA-induced EC migration and proliferation--key steps in vascular repair. Isolated EPC^miR-126^-exosomes, enriched with miR-126 (**Suppl Fig. 3B**), effectively promoted angiogenesis (**Fig. 5**). These exosomes facilitate cell-to-cell communication by transferring miRNAs to ECs, thereby bypassing the direct circulation of miRNAs, which would otherwise be susceptible to RNase degradation. This suggests that miR-126 is stable *in vivo* and that EPC^miR-126^ exosomes amplify the effects of EPCs in ischemic tissue. Although independent injections of EPC^miR-126^ exosomes as a cell-free treatment did not yield positive results-likely due to factors such as dosing and timing. Previous studies shown protective effects of EPC-derived exosomes in diabetic ischemic stroke models and neuroprotection under hypoxic conditions^45^, ^46^. Further investigation is needed to fully understand how EPC^miR-126^-exosomes and other secreted factors contribute to neurovascular repair.

Hypertension and stroke also activated cerebral vascular ECs, which increase the migration of inflammatory leukocytes from the systemic circulation into the central nervous system, thereby disrupting BBB integrity^47^, ^48^. Our study demonstrates that EPC^miR-126^ not only promotes angiogenesis but also exerts an anti-inflammatory effect on cerebral vascular endothelium. EPC^miR-126^ reduced leukocytes infiltration and preserved BBB function following stroke in SHR (**Fig. 4**). We also explored the anti-inflammatory potential of EPC^miR-126^ exosomes under stress conditions, showing that their effects could be pharmacologically suppressed by GW4869 (**Fig. 6**). Our data suggest that EPC^miR-126^ exosomes reduce endothelial inflammation by facilitating intercellular communication. These findings are consistent with previous studies that showed that miR-126 is associated with downregulation of ICAM-1 in clinical settings and decreases in pro-inflammatory cytokines, improving BBB integrity during ischemic stroke in mice^16^, ^50^.

EPC^miR-126^ presents a promising approach to overcome the limitations of traditional EPC therapies following ischemic stroke, such as poor retention, viability, and limited efficacy in the ischemic microenvironment. Exosomes derived from EPC^miR-126^ transfer miRNAs to ECs, mediating cell-to-cell communication and improving effectiveness of stem cell based therapies^41^.

However, the exert mechanism of EPC^miR-126^ in tissue repair --including direct cell replacement, secretion of trophic factors, and modulation of endogenous EPC function remain to be fully elucidated. Additionally, the use of young male SHR rats with detectable hypertension in this proof-of-concept study may limit the generalizability of these findings. Future studies should explore the effects of different cell doses, multiple treatments, and the involvement of other therapeutic factors.

In conclusion, our finding demonstrated that EPC^miR-126^ mediates ischemic tissue repair through enhanced angiogenesis and BBB protection. EPC^miR-126^-exosome serves as a crucial therapeutic agent for delivering miR-126 via paracrine secretion, promoting functional recovery in ischemic stroke. These results suggest that EPC^miR-126^ could be a promising therapeutic strategy for stroke patients, particularly those with hypertension, to enhance vascular repair and functional recovery.

## Author contributions

Study conception and design: JH, BZ, PG. Acquisition and analysis of data: DL, XL, DC, JH, MF, MQ, YS. Drafting a significant portion of the manuscript or figures: JH, DL, BZ, DC, G-YY.

## Acknowledgment

This study is supported by the National Natural Science Foundation of China (81400968 J.H., 81870311 J.H.) and the grants from the Shanghai Municipal Health and Family Planning Commission scientific research project (20154Y0130).

**Abbreviations**
BBB: blood brain barrier
BrdU: Bromodeoxyuridine
ECs: endothelial cells
EPCs: endothelial progenitor cells
EPC^miR-126^: EPC with overexpressed miR-126
MCAO: middle cerebral artery occlusion
miR-126: microRNA-126-3p
MPO: myeloperoxidase
SHRs: spontaneously hypertensive rats
OGD: oxygen-glucose deprivation
RT-PCR: real-time PCR

---

Suppl Fig. 1. **Phenotypic characterization of EPCs derived from human umbilical cord blood and mimics of miR-126 transfection. A**) Morphology of EPC cultures at different time points and different passages after plating. **B**) Flow cytometry identified cell surface markers of EPCs. Three independent experiments were performed. **C**) Immunofluorescent staining showed that EPCs were positive for KDR and CD34. **D**) Mimic miR-126 cy3 were taken up by EPCs. **E**) Relative survival rate treated with CCK-8 in EPC or EPC^miR-126^ under SHR stroke plasma treatment or OGD condition.

Suppl Fig. 2. **LV-miR-126 virus transfected EPC *in vitro*. A**) Clone of pLV-miR-126-IRES-GFP plasmid by inserting miR-126 sequence in pLV-IRES-GFP plasmid, then transfected 293 T cells with pVSVG and pDelta plasmid to package LV-miR-126 and LV-GFP virus. **B**) LV-miR-126 transfected 293T cells in fluorescent field (left), bright field (middle), merge (right). **C**) RT-PCR analysis to detect miR-126 expression in EPC^miR-126^, EPC^GFP^ and EPCs. ***, *P*<0.001

Suppl Fig. 3. **Blood pressure and body weight of SHR after MCAO. A**) Blood pressure of different treated SHR groups after MCAO. Data are mean±SEM. N=12–16 per group. **B**) Body weight of different treated SHR groups after MCAO. Data are mean±SEM. N=12–16 per group.

Suppl Fig. 4. **MiR-126 transfer through exosomes. A)** Western blot analysis show expression of exosome markers. **B**) RT-PCR results show relative miR-126 expression in exosomes from EPC^miR-126^, EPC^GFP^ and EPC. ***, *P*<0.001 **C**) Schematic diagram show EPC^miR-126^ derived exosomes could transfer into HUVECs. D) Schematic diagram shows the intercellular miR-126 trafficking between EPC^miR-126^ and HUVECs by exosomes.

Suppl Fig. 5. **IPA show miR-126 related angiogenesis candidate target genes in the cells A**) IPA predicted the miR-126 target genes in the cell nucleus cytoplasm membrane and extracellular space. **B**) Selected angiogenesis related candidate target genes of miR-126.

Suppl Table 1. Key antibodies

Suppl Table 2. Primer design

Suppl Table 3. Number of animals in accomplished experiments

